# ALC1 Finds a New Foothold on the Nucleosome’s Super-Groove

**DOI:** 10.1101/2025.11.10.687450

**Authors:** Hannah R. Bridges, Luka Bacic, Sebastian Deindl, Guillaume Gaullier

## Abstract

Nucleosomes act as recognition platforms for chromatin-binding factors that coordinate genome maintenance. A distinct structural feature of the nucleosome is the alignment of the DNA major and minor grooves across both gyres, forming continuous major and minor super-grooves. Despite this prominent feature, only one example of a naturally occurring protein binding to a super-groove is documented at the structural level to date. The chromatin remodeler Amplified in Liver Cancer 1 (ALC1) is a key component of the DNA damage response and a promising therapeutic target in cancer. Through extensive classification of our cryo-electron microscopy data, we identified a previously unresolved ALC1-nucleosome complex characterized by a loosely bound conformation of ALC1. This conformation represents an intermediate between the auto-inhibited and active states, and provides new structural insights into the conformational transitions that regulate ALC1 activity. In this intermediate state, the linker of ALC1 engages the nucleosome’s minor DNA super-groove across both gyres in a recognition mode observed for the first time in a naturally occurring protein.

**Synopsis:** Re-analysis of a publicly available cryo-EM dataset identifies an intermediate state of the oncogenic chromatin remodeler ALC1, revealing the conformational change leading to its activation upon binding to a PARylated nucleosome. The remodeler recognizes the nucleosome’s super-groove.

## 1. Introduction

### 1.1. Molecular recognition of nucleosomes

Eukaryotes package their genomes in a protein-DNA complex called chromatin. The repeating unit of chromatin is the nucleosome (Kornberg, 1974), composed of two copies each of histones H3, H4, H2A and H2B assembled as a histone octamer, around which DNA wraps as almost two turns of a left-handed super-helix (Figure 1a, b) (Luger *et al*., 1997). In addition to packaging the DNA, nucleosomes provide a recognition platform for chromatin-binding factors involved in a variety of genome maintenance processes. These proteins recognize specific epitopes on the nucleosome: the H2A-H2B acidic patch (Figure 1c), the N-terminal tail of H4, or specific post-translational modification (PTM) patterns on histone tails (Zhou *et al*., 2019; McGinty & Tan, 2021). These epitopes are often recognized in combination by chromatin-binding factors harboring multiple “reader domains” within multi-domain proteins or in multi-subunit complexes.

**Figure 1.**
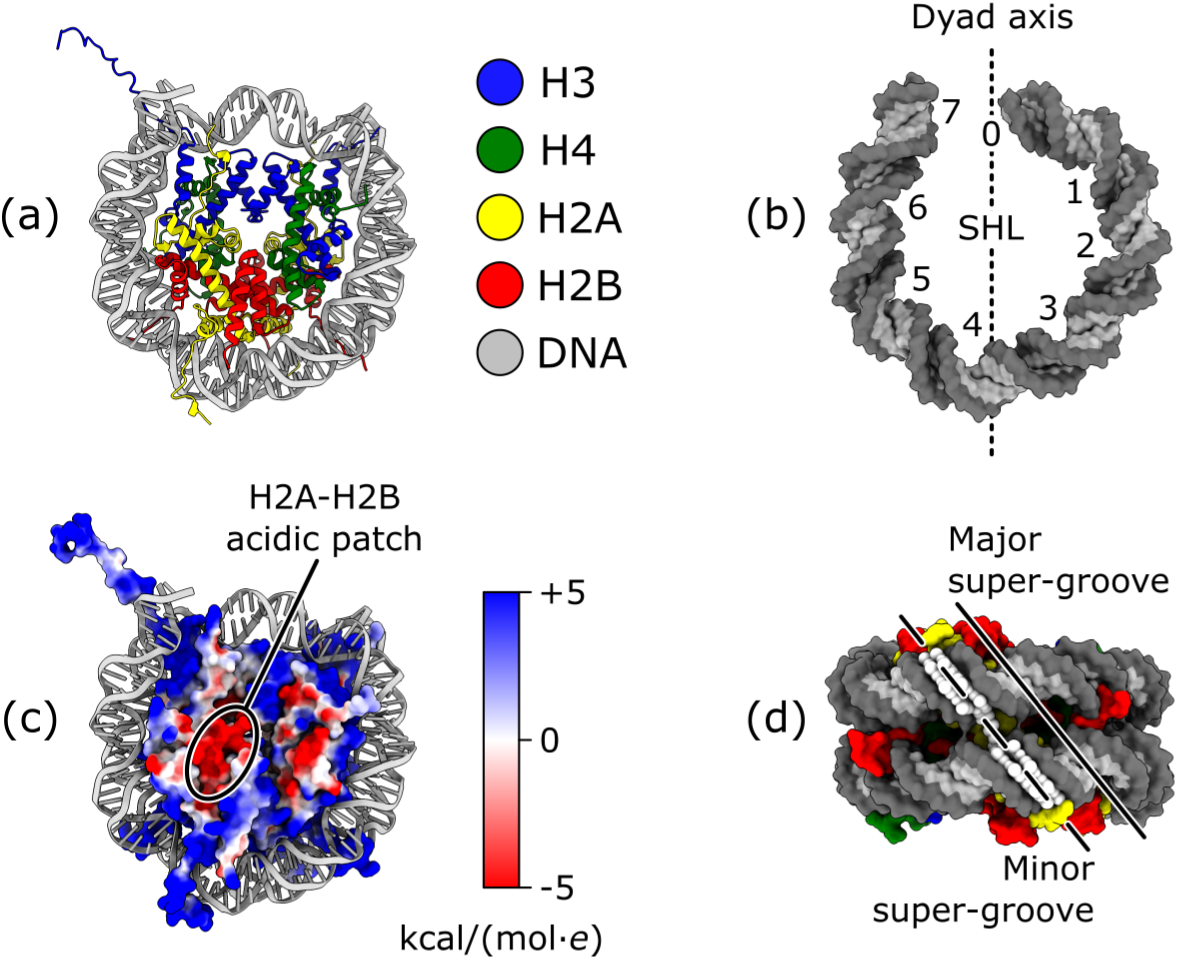
Structure of the nucleosome. (a) Overview of the nucleosome (disc view). H3 is colored in blue, H4 in green, H2A in yellow, H2B in red, and DNA in grey. From PDB 1AOI. (b) Super-helical locations (SHL) are defined as the points where the major groove faces inward, toward the histone octamer. The base pair at the center of the histone octamer footprint on the DNA is called the dyad and numbered bp 0 / SHL 0 by convention. Each gyre has 7 SHLs, numbered positively on the front gyre and negatively on the back gyre (SHLs with same numbers but opposite signs refer to equivalent locations related by the pseudo 2-fold symmetry of the dyad axis, and a fractional SHL like 1.5 refers to the minor groove between two SHLs, in this example between SHLs 1 and 2). Only the front gyre is displayed here for clarity. From PDB 1AOI. (c) Electrostatic potential of the histone octamer. The H2A-H2B acidic patch is labeled. From PDB 1AOI. (d) The nucleosome’s major and minor super-grooves (gyres view). One of the major super-grooves is indicated by a continuous line (SHL 4/-5). One of the minor super-grooves is indicated by a dashed line (SHL 4.5/-4.5). Super-grooves can be unambiguously referred to by a pair of positive and negative SHLs. The synthetic minor super-groove binder is shown as white spheres. The nucleosome is shown as a surface with the same color code as in panel (a), with the DNA backbone in darker grey. From PDB 1S32.

In addition to these canonical epitopes, non-canonical DNA structures like abasic sites, modified bases, pyrimidine dimers, nicks or double-strand breaks are recognized by DNA damage sensor proteins (Matsumoto *et al*., 2019; Gaullier *et al*., 2020; Bilokapic *et al*., 2020; Weaver *et al*., 2022; Ren *et al*., 2024). Furthermore, pioneer transcription factors recognize specific DNA sequences exposed on the outer surface of the nucleosome (Makowski *et al*., 2020; Luzete-Monteiro & Zaret, 2022).

One notable structural feature of the nucleosome is that the major and minor grooves of the DNA perfectly align across the two gyres, forming what has been termed a super-groove spanning both gyres. It was demonstrated that dimerized synthetic DNA minor groove binders can specifically recognize DNA sequences exposed in the super-groove (Figure 1d) (Edayathumangalam *et al*., 2004). Some dimeric pioneer transcription factors are suspected to target the super-groove, although there has been no direct evidence to date (Zhu *et al*., 2018; Makowski *et al*., 2020). Haspin, a histone H3 kinase involved in mitosis, has been demonstrated to bind to the cavity between the two gyres at the major super-groove of super-helical locations (SHL) 5.5/-2.5 (Hicks *et al*., 2025). With this single example of recognition of the super-groove by a naturally occurring protein described at the structural level to date, several questions remain unanswered. Is recognition of the super-groove wide-spread among chromatin-binding proteins, or restricted to certain families? Do these proteins recognize the major or minor super-groove? Which structural elements recognize super-grooves? Is recognition of the super-groove the single contributor to affinity for nucleosomes, or is it part of complex recognition modes targeting multiple nucleosomal epitopes?

### 1.2. The chromatin remodeler ALC1 in the DNA damage response

Packaging of the genome into chromatin constitutes a barrier to cellular processes that need physical access to the DNA, such as transcription factor binding and DNA damage recognition and repair. To facilitate these processes, eukaryotes have evolved a family of proteins termed ATP-dependent chromatin remodelers (Clapier & Cairns, 2009; Clapier *et al*., 2017). Remodelers can manipulate nucleosomes in a variety of ways, most prominently by performing histone exchange (to install or remove histone variants) or nucleosome translocation by sliding the histone octamer along the DNA (Narlikar *et al*., 2013; Bowman & Deindl, 2019). Like other chromatin-binding factors, remodelers follow general principles of molecular recognition of the nucleosome: they recognize nucleosomal DNA at specific SHL (mainly SHL +/- 2), the N-terminal tail of H4, and often the acidic patch (Markert & Luger, 2020). Unlike proteins involved purely in recognition and binding, chromatin remodelers undergo large conformational changes during their ATP hydrolysis cycle. This dynamic nature makes them challenging targets for structure determination: near-atomic resolution structures of remodeler-nucleosome complexes have only become available in the past decade thanks to advances of the cryo-EM “resolution revolution” (Liu *et al*., 2017; Farnung *et al*., 2017; Ayala *et al*., 2018; Eustermann *et al*., 2018; Sundaramoorthy *et al*., 2018; Willhoft *et al*., 2018; Armache *et al*., 2019; Chittori *et al*., 2019; Li *et al*., 2019; Patel *et al*., 2019; Yan *et al*., 2019; Ye *et al*., 2019; Farnung *et al*., 2020; Han *et al*., 2020; He *et al*., 2020; Wagner *et al*., 2020; Baker *et al*., 2021; He *et al*., 2021; Markert *et al*., 2021; Nodelman *et al*., 2022; Yuan *et al*., 2022; Wu *et al*., 2023; Zhang *et al*., 2023; Jalal *et al*., 2024; Li *et al*., 2024; Louder *et al*., 2024; Osakabe *et al*., 2024; Zhang *et al*., 2024; Hu *et al*., 2025; James & Farnung, 2025; Kaur *et al*., 2025; Malik *et al*., 2025; Sia *et al*., 2025; Nodelman *et al*., 2025; Tian *et al*., 2025).

Amplified in Liver Cancer 1 (ALC1) is a 101 kDa protein comprising the N- and C-ATPase domains characteristic of the chromatin remodeler family, a 200-residue linker, and a macro domain unique to this remodeler (Figure 2a). ALC1 is a key component of the DNA damage response (Gottschalk *et al*., 2009; Ahel *et al*., 2009; Gottschalk *et al*., 2012). In this cellular pathway, DNA breaks are rapidly recognized by Poly-ADP-Ribose Polymerases 1 and 2 (PARP1, PARP2) (Langelier *et al*., 2018). These two enzymes, assisted by their co-factor Histone PARylation Factor 1 (HPF1), label nucleosomes surrounding a DNA break with poly-ADP-ribose (PAR), a polymeric PTM deposited on histone tails, primarily on Ser10 in H3 and Ser6 in H2B (Leidecker *et al*., 2016; Gibbs-Seymour *et al*., 2016; Bonfiglio *et al*., 2017). ALC1 is rapidly recruited to PARylated chromatin loci, where it catalyzes chromatin relaxation to facilitate access to downstream DNA repair factors (Sellou *et al*., 2016). This recruitment happens through a direct physical interaction between PAR chains and ALC1’s macro domain, a PAR reader domain (Karras *et al*., 2005). In absence of PAR, the macro domain interacts with the ATPase domain and maintains ALC1 in an auto-inhibited state. This dual function of the macro domain is critical: its binding to PAR chains serves both as a recruitment mechanism to sites of DNA breaks, and as an activation mechanism by displacing the macro domain from the ATPase domain, thereby releasing the remodeler’s auto-inhibition (Lehmann *et al*., 2017; Singh *et al*., 2017).

**Figure 2.**
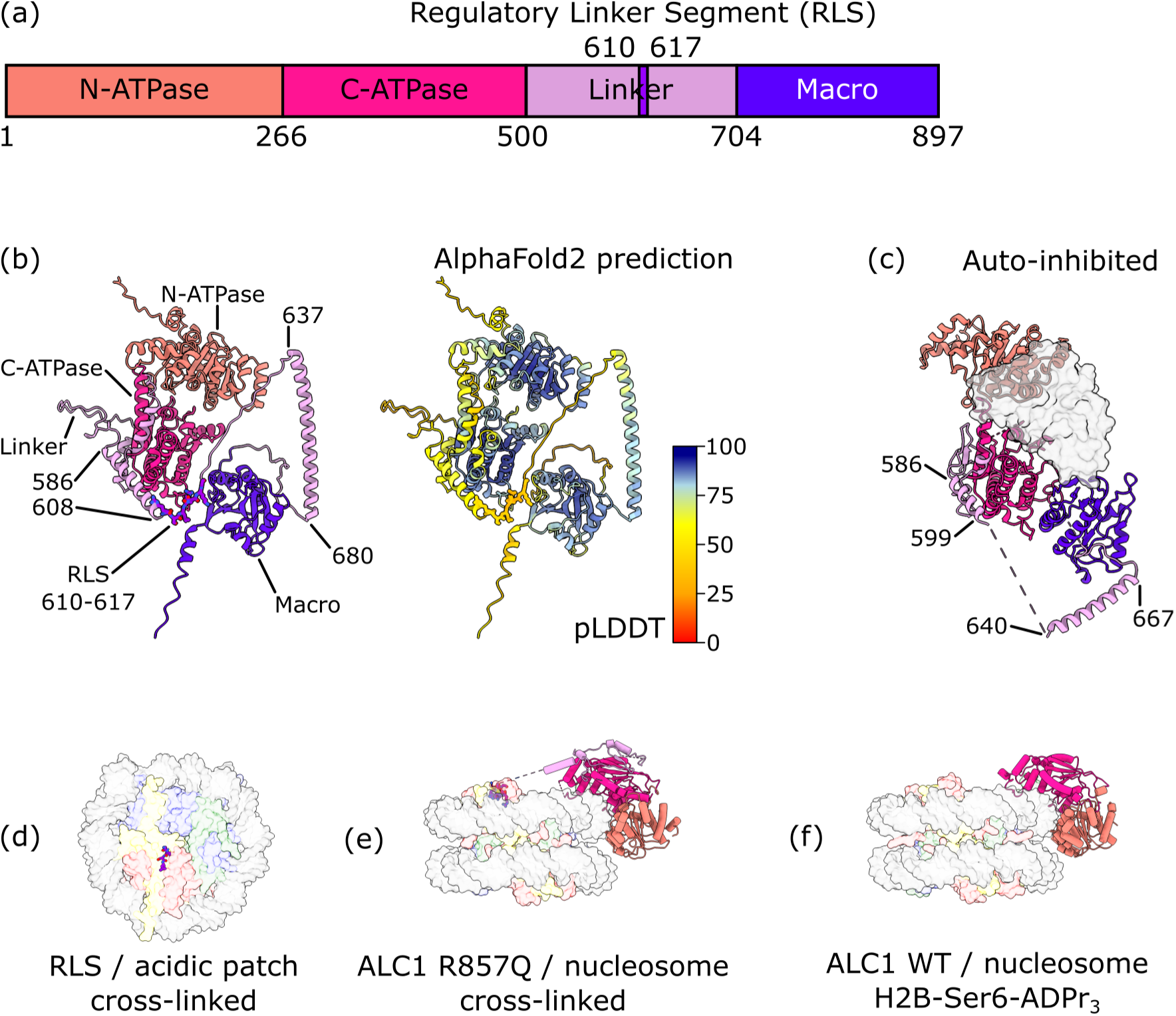
Current structural knowledge about ALC1. (a) Domain structure of ALC1. (b) AlphaFold2 prediction of full-length ALC1. Left, colored by domain as in panel (a), with domains labeled. Right, colored by pLDDT (prediction confidence). Residues of the regulatory linker segment (RLS) are displayed as sticks. From AlphaFold-DB AF-Q86WJ1-F1-v4. (c) Crystal structure of ALC1 in its auto-inhibited state, colored by domain as in (a), with the antibody used for crystallization shown as a translucent grey surface. From PDB 7EPU. (d) Structure of ALC1’s regulatory linker segment cross-linked to a nucleosome (disc view). From PDB 6ZHX. (e) Structure of ALC1 R857Q cross-linked to an unmodified nucleosome (gyres view). From PDB 7ENN. (f) Structure of ALC1 bound to a nucleosome site-specifically tri-ADP-ribosylated on H2B-Ser6 (gyres view). From PDB 8B0A. In panels (d), (e) and (f), ALC1 is colored by domain as in panel (a) and the nucleosome is colored as in Figure 1a and shown as a translucent surface for clarity.

### 1.3. Structural information currently available on ALC1

The dynamic nature of chromatin remodelers in general, and the domain structure of ALC1 in particular with its very long flexible linker between the ATPase and macro domains (Figure 2a), make structure determination challenging, since all techniques tend to work best with conformationally homogeneous macromolecular complexes. The structure prediction from AlphaFold2 is therefore valuable, as it helps with visualizing the segments of ALC1 that are either not captured, or only observed as disconnected fragments, in currently available experimental structures. Interestingly, the linker seems less disordered than had been previously thought: a long alpha helix is predicted just upstream of the macro domain (residues 637 to 680) with high confidence (Figure 2b), and indeed, residues 640 to 667 were experimentally confirmed as alpha-helical in the crystal structure of ALC1 in its auto-inhibited state (Figure 2c) (Wang *et al*., 2021).

We previously identified a regulatory element in the linker of ALC1, demonstrated that it binds to the nucleosome’s acidic patch through an arginine anchor residue (an interaction mode common among chromatin-binding factors), and determined a cryo-EM structure of this peptide bound and cross-linked to the nucleosome (Figure 2d) (Lehmann *et al*., 2020). We refer to this segment of ALC1 as the regulatory linker segment (RLS). AlphaFold2 also predicts a helix just upstream of the RLS (residues 586 to 608) with more than 50% confidence (Figure 2b), which is also partially observed (residues 586 to 599) in the crystal structure of the auto-inhibited state (Figure 2c) (Wang *et al*., 2021). This crystal structure was solved with the help of a single-chain variable fragment antibody to stabilize the auto-inhibited conformation and decrease the flexibility of the hinge between the two lobes of the ATPase domain. Neither this strategy nor the crystal packing constrained the more flexible region of the linker, so the RLS is not observed in this structure (Figure 2c) (Wang *et al*., 2021). In the same study, this group also determined a cryo-EM structure of the ALC1-nucleosome complex, by using the constitutively active ALC1 mutant R857Q in a 5:1 molar excess over unmodified nucleosomes, followed by cross-linking by the GraFix protocol (Kastner *et al*., 2008). This R857Q mutation, found in some cancers, is located in the macro domain, and abolishes both binding to PAR and auto-inhibition, resulting in unregulated chromatin remodeling by ALC1. The use of this mutant enabled the formation of a complex with an unmodified nucleosome, to which wild type (WT) ALC1 does not bind. Cross-linking likely stabilized and enabled the visualization of a part of the linker in contact with the C-ATPase lobe, and of the interaction between the RLS and the acidic patch (Figure 2e) (Wang *et al*., 2021). However, this approach prevented visualization of the macro domain (likely floating away only tethered by the end of the linker) and of any loosely bound intermediate states that were not within cross-linking distance and thus lost in purification. At the same time, we were also pursuing a structure of the ALC1-nucleosome complex, but chose a different approach. The new knowledge about the biochemistry of PARP1, PARP2 and HPF1 that developed at the time (Obaji *et al*., 2018; Gaullier *et al*., 2020; Bilokapic *et al*., 2020; Suskiewicz *et al*., 2020; Sun *et al*., 2021; Obaji *et al*., 2021; Langelier *et al*., 2021) encouraged us to prepare enzymatically PARylated nucleosomes and subsequently form a complex with WT ALC1. This resulted in a very heterogeneous sample, due to the heterogeneity of PAR chains in sites of attachment, in length and in degree of branching, but enabled the determination of a cryo-EM structure of the natural complex in its active state (Bacic *et al*., 2021). Analysis of this cryo-EM dataset with CryoDRGN (Zhong *et al*., 2021) also hinted at the presence of many more conformational states of ALC1, some clearly more loosely bound to the nucleosome, but time constraints and the image processing tools available at the time prevented us from further analyzing this complicated heterogeneity. We later used a chemoenzymatic method to produce nucleosomes homogeneously tri-ADP-ribosylated at a single site, Ser6 on a single copy of H2B (Mohapatra *et al*., 2021). This approach produced a much more homogeneous ALC1-nucleosome complex and yielded a cryo-EM structure of the active state of ALC1 at higher resolution, but in which the linker and macro domain were still not visualized (Figure 2f) (Bacic *et al*., 2024).

Overall, the structural information on ALC1 is therefore still partial and fragmented. Notably, the macro domain has never been observed in the context of an ALC1-nucleosome complex. Due to this gap in structural information, the conformational change caused by the combined recruitment and activation of ALC1 by PARylated nucleosomes has remained unclear.

Here, we revisit our heterogeneous cryo-EM dataset publicly available as EMPIAR-10739 (Bacic *et al*., 2021). We present the structure of a complex with ALC1 loosely bound to the nucleosome, representing an intermediate between the auto-inhibited and active states of the remodeler, and in which all domains of ALC1 can be assigned for the first time in the context of a nucleosome-bound structure. This intermediate state of ALC1 appears to probe a minor super-groove, a recognition mode observed in a naturally occurring protein for the first time.

## 2. Results

### 2.1. Micrograph and particle curation

The cryo-EM dataset of ALC1 bound to an enzymatically PARylated nucleosome, publicly available as EMPIAR-10739 (Bacic *et al*., 2021), was re-analyzed with the initial goal to write an educational case study for the CryoSPARC Guide (https://guide.cryosparc.com). This new analysis was also motivated by recent new developments in CryoSPARC, since the initial analysis had proven difficult in large part due to heterogeneity and to less visual feedback provided by the image processing tools available at the time. Notably, only a stringent particle picking strategy using the micrograph denoiser from Topaz (Bepler *et al*., 2020) to aid manual picking of a training set, followed by training and picking with Topaz (Bepler *et al*., 2019), had yielded reconstructions with density recognizable as ALC1.

During this re-analysis, careful curation of the micrographs was performed with two new tools in CryoSPARC: the Micrograph Denoiser (https://guide.cryosparc.com/processing-data/all-job-types-in-cryosparc/exposure-curation/job-micrograph-denoiser-beta), and the Micrograph Junk Detector (https://guide.cryosparc.com/processing-data/all-job-types-in-cryosparc/exposure-curation/job-micrograph-junk-detector-beta). A two-pass particle picking strategy was applied on the denoised micrographs: blob picking; particle curation by 2D classification, *ab initio* reconstruction and heterogeneous refinement; and template picking. The selected 2D classes included recognizable nucleosomes with high-resolution features, irrespective of whether or not they showed signal attributable to ALC1 outside of the nucleosome. This was to avoid the risk of introducing an orientation bias in viewing directions where the extra signal produces more contrast. These procedures are described in detail in sections 6.1.1 Pre-processing and 6.1.2 Particle picking and cleanup, and summarized in Figure 3 and Figure 4. Particle curation resulted in a set of particles that gave a high-resolution nucleosome reconstruction that did not appear to have density recognizable as ALC1. The lack of ALC1 density at this stage could be explained by a combination of the nucleosome itself dominating particle image alignment, sub-stoichiometric ALC1 binding, and/or a distribution of ALC1 bound at different locations and/or with different conformations that were averaged out in the reconstruction.

**Figure 3.**
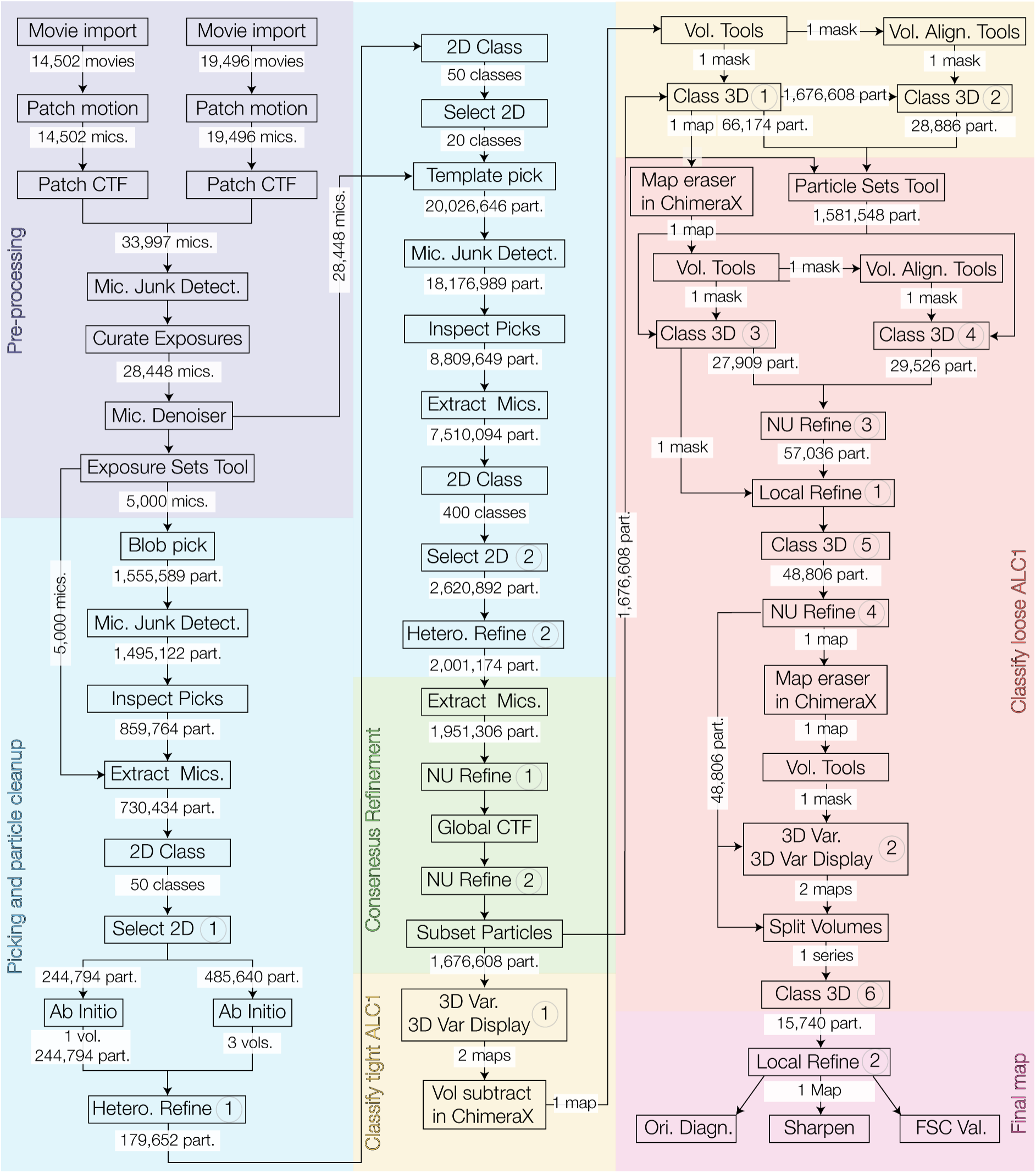
Flow chart of the image processing workflow. The steps of each distinct stage are highlighted in different colors: pre-processing in purple, particle picking and curation in blue, consensus reconstruction in green, classification of the complexes with tightly bound ALC1 in yellow, classification of the complexes with loosely bound ALC1 in red, and final refinement of the complex with loosely bound ALC1 in pink. For all steps with a circled number, key visuals are presented in Figure 4, and details are provided in section 6.1 Cryo-EM data processing.

**Figure 4.**
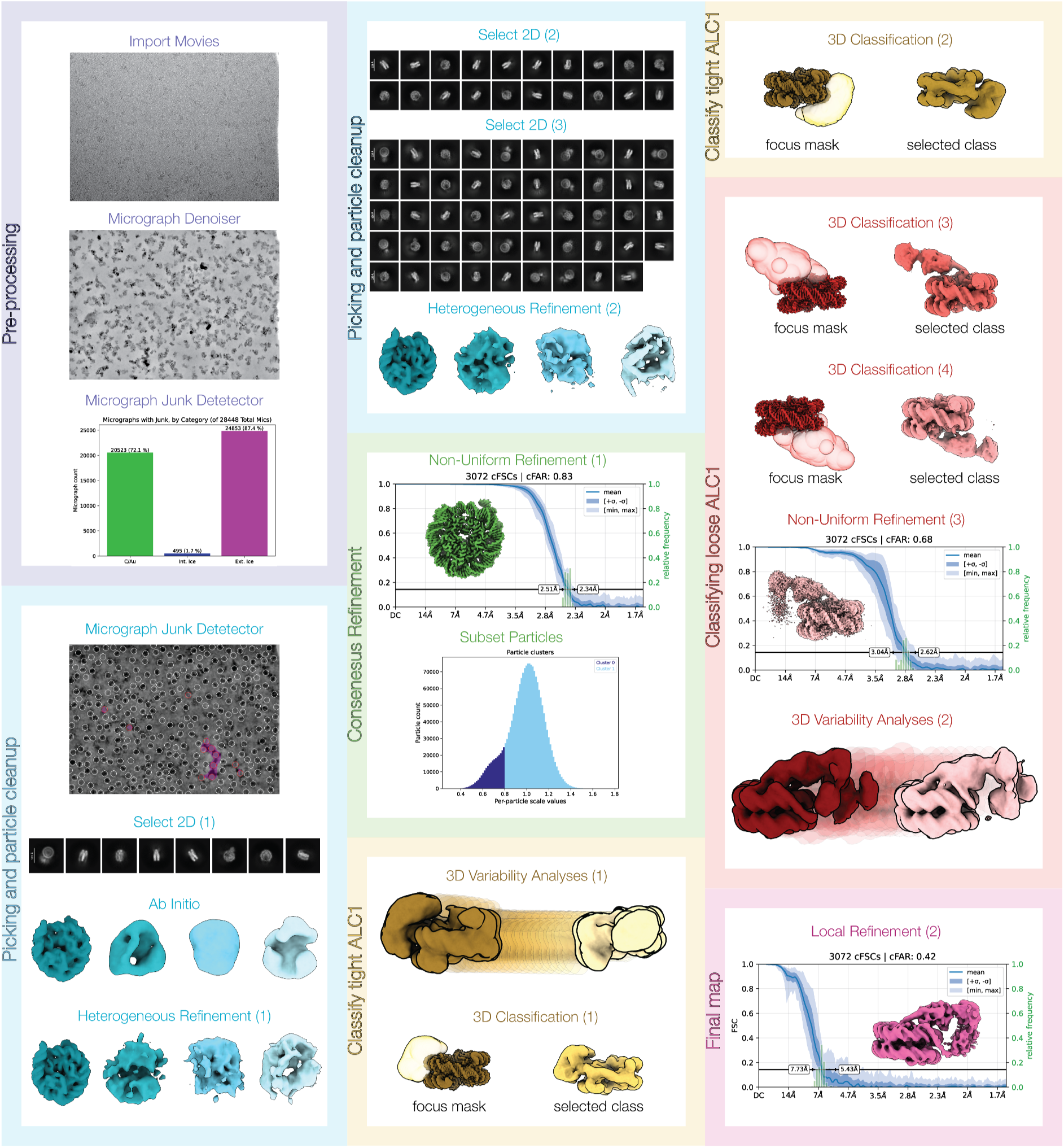
Visuals of the key processing steps from. Figure 3. The background follows the same color code as in Figure 3. The numbers in parentheses refer to the circled numbers in Figure 3, and details about these are provided in section 6.1 Cryo-EM data processing.

### 2.2. Extensive 3D classification identifies a complex with a loosely bound ALC1

The tightly bound conformation of ALC1 was first reported in (Bacic *et al*., 2021), and was found following extensive 3D classification of the particles. Classification of tightly-bound ALC1 was also performed as part of reprocessing this EMPIAR-10739 dataset as a case study for the CryoSPARC Guide (https://guide.cryosparc.com/processing-data/tutorials-and-case-studies/case-study-end-to-end-and-exploratory-processing-of-a-motor-bound-nucleosome-empiar-10739). This procedure involved 3D variability analysis (3DVA) on the particles from the consensus refinement, and masked 3D classifications, as detailed in section 6.1.4 Classification of tightly-bound ALC1. We will not further describe this tightly bound complex here.

The initial analysis of this dataset followed three strategies. First, via multiple rounds of hierarchical 3D classification in RELION, the alignments may have degraded as the set of particles was iteratively split into more and more classes, each one populated by fewer particles. Perhaps not surprisingly, this strategy did not produce any interpretable reconstruction. The second strategy used 3DVA in cluster mode (CryoSPARC version 3.2) and isolated a set of particles corresponding to the tightly bound state. The third strategy used CryoDRGN, recently published at the time, and produced reconstructions of multiple conformational states of ALC1 all consistent with previous biochemical information (EMD-13070). But none of the maps from CryoDRGN had sufficient resolution to rigid-body fit the domains of ALC1 confidently, and the distribution of latent space coordinates of the particles did not reveal discrete clusters that could be separately subjected to conventional refinement and reconstruction.

In this re-analysis, a combination of 3DVA (Punjani & Fleet, 2021) and extensive 3D classification (with no alignment) using wide masks encompassing the space occupied by ALC1 eventually produced a 3D reconstruction of a loosely-bound state of ALC1. The procedure is detailed in section 6.1.5 Classification of loosely-bound ALC1 and summarized in Figure 3 and Figure 4. It was critical to consider the fact that ALC1 could bind on either side of the nucleosome due to the heterogeneous PARylation pattern on the nucleosome. Therefore, 3D classification needed to be performed for both sides separately. Note that this was not performed as a single classification with C2 symmetry expansion because we wanted to first ensure that the asymmetric DNA sequence had not visibly influenced the binding of ALC1, and that its conformation was the same on both sides. The resulting reconstructions were realigned to match ALC1 on a single side. Although this will have contributed to blurring the asymmetric DNA sequence, it improved the quality of the density for ALC1.

### 2.3. The loosely bound ALC1 is an intermediate between the auto-inhibited and active states

This extensive curation and classification procedure identified an ALC1-nucleosome complex with a loosely bound ALC1, in which all domains of ALC1 are finally visualized in the context of a nucleosome complex (Figure 5a). The final classified particle set contains 15,740 particles, which constitutes only ∼1% of the curated particles in the consensus refinement. We note that the map quality for the N-ATPase lobe and macro domain is relatively poor compared to the nucleosome and C-ATPase lobe, as reflected by the local resolution estimates (Figure S3). This likely indicates unresolved heterogeneity, but attempts to further sub-classify did not improve the map quality in these regions, probably limited by the small number of particles remaining. The map still enabled unambiguous rigid-body fitting of all domains (for details, see section 6.2 Model building and refinement), allowing interpretation of their relative orientations and interactions at the secondary structure level.

**Figure 5.**
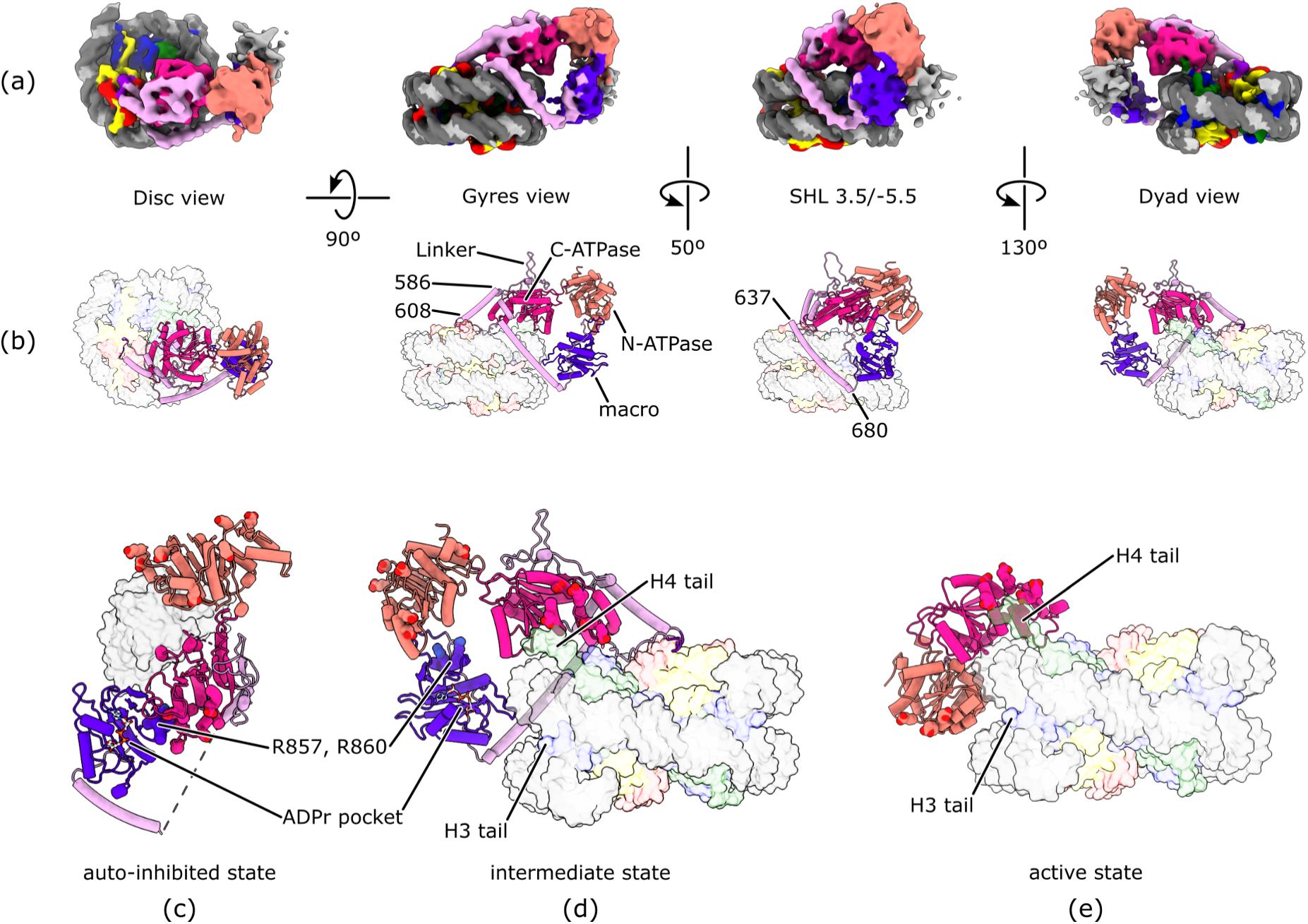
Structure of an activation intermediate of ALC1 loosely bound to a PARylated nucleosome. (a) Cryo-EM map colored by chain assignment, following the color codes from Figure 1a for the nucleosome and Figure 2a for ALC1. (b) Atomic model, shown in the same orientations and with the same color code as the map shown in panel (a). (c) Structure of the auto-inhibited state. The antibody used for crystallization is shown as a translucent grey surface. From PDB 7EPU. (d) Structure of the intermediate state (dyad view). From this study. (e) Structure of the active state. From PDB 8B0A. In panels (c), (d) and (e), the ATPase residues mutated in the study by (Lehmann *et al*., 2017) are shown as spheres. In panels (c) and (d), the macro domain residues R857 and R860, found mutated in cancer, are shown as spheres and labeled, and the ADPr-binding pocket of the macro domain is indicated by an ADPr molecule (shown as balls-and-sticks) modeled by structural superimposition of the structure of the Af1521/ADPr complex from PDB 2BFQ (the Af1521 protein is not shown for clarity). In panels (b), (d) and (e), the nucleosome is shown as a translucent surface for clarity.

This structure presents unique features not observed in the structures of the auto-inhibited and active states (Figure 5b). The C-ATPase lobe interacts with the H4 tail and with the DNA, but not quite at SHL 2 where it eventually settles in the active state. This is consistent with our previous analysis with CryoDRGN that had revealed that ALC1 samples the nucleosome at a range of locations between SHL 1 and 3 (Bacic *et al*., 2021). The RLS interacts with the acidic patch, likely more stably than in the active state in which the RLS was only detected in a cross-linked structure (Figure 2e). The N-ATPase lobe does not yet interact with the DNA, and is instead still interacting with the macro domain. The macro domain interacts with both the N-ATPase lobe and the DNA. The macro domain most likely also interacts with the PAR chains that are too heterogeneous and flexible to be visualized. This is supported by the observations that the ADPr-binding pocket of the macro domain is exposed in the auto-inhibited state (Figure 5c), and points towards the tail of H3 (where H3-S10, the main site of histone PARylation, is located) in this new structure (Figure 5d). Because the ATPase domain is not clamped on the DNA at SHL 2 (compare Figure 5d and e), this structure does not represent a catalytically active state. The conformation of ALC1 observed here is however markedly different from its auto-inhibited state: the macro domain interacts with the N-ATPase lobe instead of the C-ATPase lobe (compare Figure 5c and d). This is consistent with previous evidence from small-angle X-ray scattering and cross-linking mass spectrometry showing that the macro domain could interact with both ATPase lobes (Lehmann *et al*., 2017). The sample contained ADP-BeF_3_, an ATP analogue that traps remodelers in their tightly bound conformation on nucleosomes. Assuming the particles that entered this tightly bound conformation could not dissociate, this new structure must therefore represent an intermediate state traversed by ALC1 between its initial recruitment to PARylated nucleosomes and its eventual activation.

### 2.4. ALC1 recognizes the nucleosome’s super-groove

This new structure reveals that the linker of ALC1 is in fact much less disordered than previously thought. Residues 507 to 519 and 522 to 527 form two short alpha helices that rest on the surface of the C-ATPase lobe. Residues 586 to 608 form an alpha helix leading into the RLS, likely providing enough rigidity in the linker to restrict the conformational freedom of the RLS and help point it at the acidic patch. Residues 528 to 585 don’t form regular secondary structures, but seem to interact with the C-ATPase lobe. This segment of the linker might provide some flexibility to balance the rigid helix 586-608, possibly facilitating the interaction between the RLS and the acidic patch by helping bridge a variable distance between the acidic patch and the C-ATPase lobe before its interactions with the H4 tail and DNA restrict its movement.

The most striking feature of the structure of this intermediate state is a long alpha helix from residues 637 to 680 of ALC1, that closely tracks the nucleosome’s minor super-groove at SHL 3.5/-5.5. This helix is rich in basic residues, with 6 arginines and 9 lysines representing together 34% of its 44 residues. These basic residues are distributed along the entire length of the helix (Figure 6a), point in all directions around the helical axis (Figure 6b), and those facing the DNA are conserved among ALC1 orthologues from vertebrates (Figure 6b, c). Further support for the function of this helix in binding to nucleosomal DNA comes from our previous study, in which we observed that a macro domain construct starting at residue 613 (therefore encompassing this entire helix) shifted unmodified nucleosomes in electrophoretic mobility shift assays (Bacic *et al*., 2021). From here on, we refer to this structural element of ALC1 as the super-groove recognition helix (SGRH).

**Figure 6.**
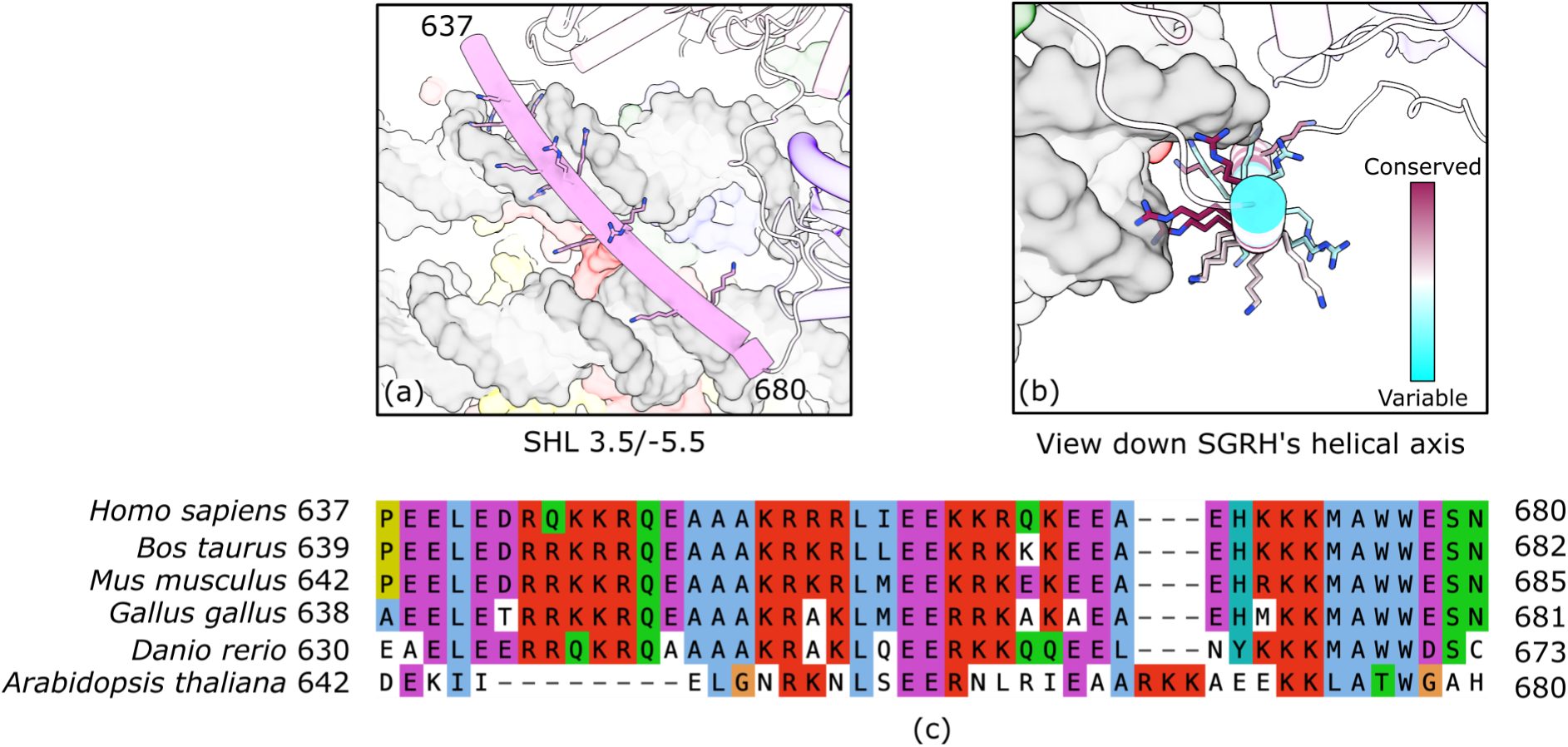
The super-groove recognition helix of ALC1. (a) View of the minor super-groove at SHL 3.5/-5.5, with the basic residues of the SGRH shown as sticks, and the residue numbers at the start and end of the helix indicated. The color code is the same as in Figure 5. The rest of ALC1 and the nucleosome are translucent for clarity. (b) View of the SGRH looking down the helical axis, with the SGRH colored by conservation. (c) Multiple sequence alignment of ALC1 orthologues verified by structure predictions to have a similar domain architecture as human ALC1, including a putative SGRH.

## 3. Discussion

### 3.1. On finding the unexpected

By revisiting the publicly available cryo-EM dataset from our previous study, we discovered an ALC1-nucleosome complex in which ALC1 is loosely bound, representing an intermediate between its auto-inhibited and active states. Based on our experience working on ALC1, we believe that no biochemical manipulation would have enabled us to trap and enrich this intermediate state. Therefore, biochemical reconstitution mimicking an *in vivo* situation (however complex) as closely as possible, paired to extensive classification during image processing, is a strategy that could be more commonly employed when working on chromatin complexes or other heterogeneous systems.

### 3.2. Function of the super-groove recognition helix in the activation of ALC1

A hallmark of this newly visualized intermediate state of ALC1 is that a segment of its linker probes the nucleosome’s super-groove at SHL 3.5/-5.5 with a 44-residue long, basic helix we termed the super-groove recognition helix (SGRH). This helix could have at least two, non-mutually exclusive functions.

The SGRH could participate in recognition of the nucleosome, in addition to other structural elements of ALC1 interacting with other nucleosomal epitopes: the macro domain interacting with PAR chains, the RLS interacting with the acidic patch, and the ATPase domain interacting with the H4 tail and the DNA at SHL 2. It is unclear why ALC1 would need such a high redundancy in recognition elements, since simultaneously probing the H4 tail and the acidic patch already guarantees that the bound substrate is a nucleosome, and since the directionality of nucleosome sliding is established by the PARylation pattern (Bacic *et al*., 2024). Basic residues pointing in all directions around the SGRH’s helical axis may help it interact with nucleosomal DNA throughout the large conformational change undergone by ALC1 during its activation.

The SGRH could also be an integral part of the activation mechanism, for example by acting as a mechanical lever to help dissociate the macro domain from the ATPase domain. This hypothesis is supported by the fact that residues 640 to 667 of the linker are already folded as a helix in the auto-inhibited state, while residues 668 to 680 are not (Figure 2c). This 668-680 segment likely folds upon interaction with nucleosomal DNA, completing the SGRH. This transition from coil to helix would stiffen this last segment, which might help pull the macro domain off the ATPase domain. The following sequence of events for the activation of ALC1 is compatible with all functional and structural evidence available: (1) binding of the macro domain to PAR chains recruits ALC1 to a nucleosome; (2) the H4 tail out-competes the macro domain for binding to the C-ATPase lobe, the SGRH binds to the minor super-groove at SHL 3.5/-5.5, and these interactions pull the C-ATPase lobe and macro domain apart, forcing them to dissociate; (3) the C-ATPase lobe finds its correct orientation because it gets anchored by three points: its interactions with the H4 tail and the DNA, and the interaction of the RLS with the acidic patch; (4) the macro domain “rolls over” to interact with the N-ATPase lobe instead; (5) these two domains eventually dissociate, allowing the N-ATPase lobe to close the ATPase domain’s clamp on the DNA at SHL 2 (a hallmark of the active state). Of note, CHD1, the closest paralogue of ALC1, undergoes similar conformational transitions involving interactions between several of its domains and regulatory structural elements (James & Farnung, 2025).

These hypotheses on the activation mechanism of ALC1 will require experimental testing in future functional studies.

### 3.3. Recognition of the nucleosome’s super-groove

The structure we report here is the second example of recognition of the nucleosome’s super-groove by a naturally occurring protein, after the structure of a Haspin-nucleosome complex (Hicks *et al*., 2025). Interestingly, the binding modes of Haspin and ALC1 are very different: Haspin inserts its kinase domain in the cavity formed by the two DNA gyres at the major super-groove, while ALC1 features a basic alpha helix long enough to span an entire minor super-groove across the two gyres. These two different binding modes raise new questions: can super-groove recognition be achieved by more diverse structural elements? Is recognition of the super-groove wide-spread among chromatin-binding factors? Are super-grooves always recognized independently of their DNA sequence?

The example of ALC1, with its SGRH located just upstream of its macro domain, suggests a more general class of structural elements targeting the super-groove: a basic helix approximately 40 residues in length, followed by a short linker connecting it to any histone PTM reader domain. Because histone tails protrude out of the nucleosome from between the two gyres (for H3 and H2B) or from the outer edge (for H4 and H2A), interaction of a histone PTM reader domain with its target on a histone tail is likely to bring a neighboring structural element in close proximity to the nucleosomal DNA. Such arrangements of structural elements may be searched systematically by bioinformatic means.

### 3.4. On the importance of making raw data publicly available

We note and emphasize that the findings presented here would most likely not have been made, had we not deposited the dataset from our 2021 study in EMPIAR (Iudin *et al*., 2023). We hope this report will encourage other structural biologists to deposit the raw data supporting their structural models into public databases, and that it will also encourage downstream users of public datasets to reach out to the depositors when new discoveries are made. Fruitful collaborations can ensue, as we hope we have demonstrated here.

## Supporting information

ChimeraX command file to set up a session with the same color code as in the figures

Multiple sequence alignment of ALC1 orthologues used for the conservation analysis (fasta format)

## 4. Data availability

The cryo-EM dataset re-analyzed in this study is publicly available as EMPIAR-10739. The reconstruction of the ALC1-nucleosome complex in an activation intermediate state is deposited in the EMDB with accession code EMD-55533 (the map series from 3DVA (2) is deposited in the same entry as additional maps) and the corresponding atomic model is deposited in the PDB with accession code pdb_00009T4V. The reconstruction used to model the RLS with ModelAngelo is deposited in the EMDB with accession code EMD-55534.

## 5. Author contributions

HRB carried out all image processing and wrote the corresponding method section, performed initial model building with ModelAngelo, and prepared figures. LB surveyed the literature and the PDB to search for other examples of super-groove binding modes. SD funded and supervised the initial work that produced the cryo-EM dataset revisited in this study. GG performed model building and refinement, carried out the conservation analysis, prepared figures, and wrote the initial version of the manuscript. All authors discussed the results and their interpretation, and edited the manuscript.

## Acknowledgements

We thank the entire team at Structura Biotechnology Inc. that designs, develops, maintains and supports the CryoSPARC software system. HRB is an employee of Structura Biotechnology Inc. GG thanks Cecilia Blikstad for access to a GPU workstation, Jonathan W. Markert for intellectually stimulating discussions about all things chromatin, and Rebecca M. Waterhouse for her patience and support during the evenings and weekends spent working on this article.

## 6. Methods

### 6.1. Cryo-EM data processing

All processing steps were conducted using CryoSPARC version 4.7.1 (Punjani *et al*., 2017) and ChimeraX version 1.9 (Pettersen *et al*., 2021). Statistics are reported in Error: Reference source not found.

#### 6.1.1. Pre-processing

Movies from EMPIAR-10739 were imported to CryoSPARC as two separate jobs, one each for the two data collection batches comprising 14,502 and 19,496 movies for batches 1 and 2, respectively, thereby assigning the two batches to different exposure groups. In-movie motion was corrected using Patch Motion, saving the images in 16-bit floating point, followed by estimation of micrograph contrast transfer function (CTF) by Patch CTF with default settings. The two micrograph sets were then input into a single Micrograph Junk Detector job with default settings, to identify regions of junk within the images. While a high number of micrographs contained features that resembled foil hole edge (72.5%) and extrinsic ice defects (such as small ice crystals or ethane contaminants) (87.5%), the image area that these took up was relatively low at 4.9% and 3.3% respectively. Using Manually Curate Exposures, poor quality micrographs were rejected by using the criteria of max CTF fit resolution (Å):5, max Intrinsic Ice Defect Area (%):10, max Total full-frame motion distance (pixels): 20, and max Relative Ice Thickness: 1.2. This left 28,448 accepted micrographs (83.7%) that were enhanced by using the Micrograph Denoiser job using default settings, to enable higher quality particle picking.

#### 6.1.2. Particle picking and cleanup

Particle picking was performed in two stages; initial blob picking to generate 2D class averages, followed by template picking. To speed up the generation of good 2D class averages, a subset of the accepted micrographs was used initially, using Exposure Sets Tool, setting Split batch size: 5000. Blob picker was run on these 5,000 micrographs using particle diameter 150-180Å, and enabling Pick on denoised micrographs. The Micrograph Junk Detector job was used to automatically reject particle picks that were on regions of junk, and Inspect Particle Picks was used in Cluster mode with Target power score: 100. 730,434 particles were extracted with a box of 350 pixels Fourier cropped to 64, and saved in 16-bit floating point. Extracted particles were 2D classified using a maximum resolution of 10 Å, initial uncertainty factor:1, and 2D zeropad factor:1. Eight classes resembling a nucleosome were selected using Select 2D Classes (1). To further enhance the quality and diversity of nucleosome views present in the 2D class averages, remaining poor particles were removed before re-running 2D classification; 1 class Ab Initio Reconstruction was run using 20,000 of the selected particles from Select 2D, 3 class Ab Initio Reconstruction was run using 1,000 of the excluded particles from Select 2D. For this second Ab Initio job, the maximum resolution was set at 30 Å, Number of initial iterations: 20 and Number of final iterations: 30. These settings were used to rapidly generate decoy volumes. All 4 volumes from Ab Initio jobs were input to Heterogeneous Refinement (1), along with the accepted particles from Select 2D, setting Refinement box size (voxels): 64. Only one of the resulting volumes resembled a nucleosome, so the associated particles were input to 2D Classification with a maximum resolution of 10 Å, initial uncertainty factor: 1, and 2D zeropad factor: 1 and 30 classes. The best 20 classes were selected using Select 2D Classes (2) and these class averages showed a mixture of views: disc, gyres, dyad and some views titled between those (nomenclature of nucleosome views according to Zhou *et al*., 2019). Some of the class averages also showed density attached to the nucleosome that might represent ALC1, but classes that did and did not show this density were included. Template Picker was run on all 28,448 accepted micrographs using a Particle diameter of 140 Å, and enabling Pick on denoised micrographs. The Micrograph Junk Detector job was used to automatically reject particle picks that were on regions of junk, and Inspect Particle Picks was used in manual mode to set a Local power score range of 57-223, and NCC score or >0.48. 7,510,094 particles were extracted with a box of 350 pixels Fourier cropped to 64, and saved in 16-bit floating point. Extracted particles were input to 2D Classification with 400 classes, a maximum resolution of 12 Å, initial uncertainty factor: 0.5 and 2D zeropad factor: 1. The large number of classes and low uncertainty factor were used to improve separation of diverse types of junk. The best 49 classes were selected using Select 2D Classes, and input to Heterogeneous Refinement (2) using the previously generated Ab Initio volumes, and setting Refinement box size (voxels): 64.

#### 6.1.3. Consensus Refinement

To enable high resolution 3D refinement, and to re-center images after alignment in 3D, the 1,951,306 particles from the good Heterogeneous Refinement class were re-extracted with a box size of 416 pixels without Fourier cropping, and saved in 16-bit floating point. Due to the asymmetric nature of the Widom 601 DNA sequence used to assemble these nucleosomes (Lowary & Widom, 1998; Ngo *et al*., 2015), Non-Uniform (NU) Refinement (1) with the re-extracted particles, and Good Ab Initio volume with C2 symmetry and symmetry relaxation by marginalization over poses were run. Minimize over per-particle scale was enabled, and the Dynamic mask start resolution was set to 1 to effectively disable masking. The resulting map achieved a nominal FSC=0.143 resolution of 2.45 Å and cFAR score of 0.83, together indicating the likelihood of good orientational sampling of high-quality images, but the map did not show any density that might belong to ALC1. To further improve the map quality, Global CTF Refinement was run including Tilt, Trefoil, Spherical Aberration, Tetrafoil and Anisotropic Magnification. The resulting particles were input to NU Refinement (2) with the same settings as NU Refinement (1) and with the addition of enabling Optimize over per-particle defocus, Optimize per-group CTF params, Fit Spherical Aberration and Fit Tetrafoil. The resulting map achieved a nominal FSC=0.143 resolution of 2.26 Å and cFAR of 0.80. A bimodal distribution of the per-particle scale factors was observed, manifesting as a shoulder in the low-scale region. This often indicates poor quality or junk particles, so Subset Particles (https://guide.cryosparc.com/processing-data/all-job-types-in-cryosparc/particle-curation/job-subset-particles-by-statistic) with subset by per-particle scale, subsetting mode: Split by manual threshold and threshold 1: 0.8 was run to separate particles in the main peak and shoulder. Cluster 1 (the high scale cluster) contained 1,676,608 particles and was used for downstream processing.

#### 6.1.4. Classification of tightly-bound ALC1

Previous work on this dataset identified a class of nucleosomes that had a tightly bound ALC1 ATPase domain (Bacic *et al*., 2021), but the maps obtained here so far did not show density for ALC1. A generous solvent mask was generated by using Volume Tools with the volume from NU Refinement (2), Lowpass filter of 15 Å, binarization threshold of 0.025, Dilation radius (pix): 25 and selecting Type of output volume: mask. To investigate heterogeneity in the sample, 3D Variability Analysis (1) was run using 100,000 particles and the generous solvent mask. 3D Variability Display was then run in simple mode with Downsample to box size:128, and the resulting volume series viewed using ChimeraX. One of the modes showed the appearance and disappearance of density bound to the side of the nucleosome. The first and last frame volumes from this mode were used to make a difference map in ChimeraX by using the *volume subtract* command, then small blobs of the remaining density were cleaned up by using the volume eraser tool to leave a volume encompassing only the ALC1 region. This volume was uploaded and imported to CryoSPARC and used in a Volume Tools job to generate Focus mask (1) by using a lowpass filter of 15 Å, Threshold of 0.019, Dilation radius of 4 and soft padding width of 15 pixels, and selecting Type of output volume: mask. Because these nucleosomes were enzymatically PARylated, resulting in a heterogeneous pattern of PARylation, ALC1 could potentially bind to both sides of the nucleosome. Therefore, classification was performed on both sides separately. Taking advantage of the volume being aligned along the C2 symmetry axis of the nucleosome, Volume Alignment Tools was used on the imported ALC1 volume and Focus mask (1) with 3D rotation Euler angles (deg or rad): 0,0,180 deg to produce Focus mask (2). Focus mask (1) along with the generous solvent mask, were used in 3D Classification (1) of the Cluster 1 particles into 40 classes, with Initialization mode: PCA, Filter resolution (Å): 15 and Class similarity: 0.1. Most of the resulting volumes did not show clear density for ALC1, but a single class appeared to represent the tightly-bound state of ALC1, and one contained less well-ordered density in the same region but with a different shape. Focus mask (2) was not contained within the generous solvent mask, so a very generous solvent mask was created using the same settings except a Dilation radius of 100 pixels. 3D Classification (2) was performed on the Cluster 1 particles into 80 classes, using the very generous solvent mask and Focus mask (2) with Initialization mode: PCA, Filter resolution (A): 15 and Class similarity: 0.1. Most of the resulting volumes did not show clear density for ALC1, but a single class appeared to represent the tightly-bound state of ALC1. We will not further discuss processing of the tightly-bound ALC1 classes, as they represent the same conformation discussed in previous work (Bacic *et al*., 2021, 2024).

#### 6.1.5. Classification of loosely-bound ALC1

The volume observed in 3D Classification (1) that had less well-ordered density in the ALC1 region was examined in ChimeraX, the core nucleosome density was removed by using the map eraser tool, and the map was smoothed by using the *volume gaussian* command. This volume was uploaded and imported to CryoSPARC, and used in a Volume Tools job to generate Focus mask (3) by using a lowpass filter of 15 Å, Threshold of 0.05, Dilation radius of 6, and selecting Type of output volume: mask. In a similar strategy as for the tightly-bound ALC1 state, Volume Alignment Tools was used on the imported ALC1 volume and Focus mask (3) with 3D rotation Euler angles (deg or rad): 0,0,180 deg to produce Focus mask (4). As the particles with tightly-bound ALC1 were already identified in 3D Classifications (1) and (2), these particles were excluded from downstream processing by using Particle Sets Tool, and inputting Subset Particles Cluster 1 in the Particles A slot, the two particles sets corresponding to the tightly-bound ALC1 classes from Classifications (1) and (2) in Particles slot B, and setting Action: intersect. The remaining particles in A minus B (1,581,548) were then used for 3D Classification (3) into 40 classes using the very generous solvent mask, Focus mask (3) with Initialization mode: PCA, Filter resolution (Å): 5 and Class similarity: 0.1, O-EM batch size (per class): 300 and O-EM learning rate: 0.9. Classification (4) was performed with the same settings, but using Focus mask (4). In each of 3D Classifications (3) and (4), there was a single class that contained good density in the region of ALC1 comprising 27,909 and 29,529 particles, respectively. These particles were combined and input to NU Refinement (3), using the selected volume from Classification (3) as an input reference, using the same settings as for NU Refinement (2) except for using C1 symmetry. The resulting map had a nominal FSC=0.143 resolution of 2.82 Å and cFAR of 0.68 (Figure S1). The map showed density for a small region making contact with the nucleosome acidic patch that was of sufficient quality to assign this as part of ALC1 (see section 6.2 Model building and refinement), but the rest of the ALC1 density was of poor, featureless quality. The map was examined in ChimeraX, the core nucleosome density was removed by using the map eraser tool, and the map was smoothed by using the *volume gaussian* command. This volume was uploaded and imported to CryoSPARC, and used in a Volume Tools job to generate Focus mask (5) by using a lowpass filter of 15 Å, Threshold of 0.05, Dilation radius of 6, and selecting Type of output volume: mask. To try and compensate for continuous heterogeneity, Local Refinement (1) was performed using Focus mask (3) with an Initial Lowpass resolution of 8 Å, maximum alignment resolution of 5 Å, enabling Re-center rotations each iteration, Re-center shift each iteration, and Use pose/shift gaussian prior during alignment, along with Standard deviation (deg) of prior over rotation:3, and Standard deviation (Å) of prior over shifts: 2. The resulting map had a nominal FSC=0.143 resolution of 4.76 Å, and showed more continuous density outside of the nucleosome, but was still of poor quality. To remove remaining poor particles, or those that struggled to align locally, 3D Classification (5) into 10 classes was performed using the Local Refinement input, Focus mask (3) using Initialization mode: PCA, Filter resolution (Å): 5, Class similarity: 0.1, O-EM batch size (per class): 300 and O-EM learning rate: 0.9. Five of the output classes (totaling 48,806 particles) showed equivalent strong density in the masked region. To further investigate the heterogeneity that was limiting map clarity in the region of interest, a new mask was made to cover the additional region of density. The volume from Local Refinement (1) was examined in ChimeraX, the core nucleosome density was removed by using the map eraser tool, and the map was smoothed by using the *volume gaussian* command. This volume was uploaded and imported to CryoSPARC, and used in a Volume Tools job to generate Focus mask (6) by using a Threshold of 0.03, Dilation radius of 20, and selecting Type of output volume: mask. The selected particles from 3D Classification (5) were subjected to NU Refinement (4) with the same settings as NU Refinement (3) followed by 3DVA with Focus mask (6) using 5 modes, and a filter resolution of 12 Å. 3D Variability Display was then run in simple mode with Filter Resolution:6 and Downsample to box size:128, and the resulting volume series 3DVA (2) was viewed using ChimeraX. One of the modes showed the motion of a tube of density, apparently inserting in the minor DNA groove. This appeared to represent two discrete states, rather than continuous heterogeneity (Figure S2). Split Volumes was run using the 3DVA series containing the interesting motion, and the volumes for frames 0 and 19 were input to a 2-class 3D Classification (6) of the particles from NU Refinement (4) with Initialization mode: input, Filter resolution (Å): 10, O-EM batch size (per class): 200 and O-EM learning rate: 0.3. One class containing 15,740 particles displayed a long helical density inserting in the DNA minor groove.

#### 6.1.6. Final refinement, sharpening and validation

The particles from this last class were subjected to Local Refinement (2) using the same settings as for Local Refinement (1). A generous mask was made from the output volume using Volume Tools with a Lowpass Filter of 15 Å and dilation radius of 10 pixels, followed by running the FSC validation job. The global resolution at FSC=0.143 was estimated at 6.60 Å. The same mask and map were used for Local Resolution and Local filter jobs, which indicated local resolutions at FSC=0.5 of ∼5-18 within the masked region. The same mask along with the locally refined map and particles were also used in an Orientation Diagnostics job. The local refined map displayed a cFAR of 0.42 indicating the possibility of a moderate preferred orientation that was not present in the consensus refinement, and an SCF* of 0.872 (Figure S3). Finally, the locally refined map was sharpened with a B-factor of -300 Å^2^ using the Sharpen job.

### 6.2. Model building and refinement

An initial atomic model was prepared using ChimeraX version 1.10.1 (Pettersen *et al*., 2021), by placing fragments into the map manually, or by using the *matchmaker* command between segments of overlapping sequence in the different fragments, followed by rigid-body fitting of each fragment with the *fitmap* command. All ALC1 fragments were taken from the AlphaFold2 prediction (AlphaFold-DB accession code AF-Q86WJ1-F1-v4), except for a fragment built by ModelAngelo (Jamali *et al*., 2024). The following fragments were placed in this order: the entire nucleosome from PDB entry 8B0A, ALC1 residues 599-626 (acidic patch recognition element) built by ModelAngelo (using the full-length sequence information) from the map from NU Refinement (3), ALC1 residues 581-609 (linker helix) by their overlap with the previous fragment, ALC1 residues 627-874 (macro domain and super-groove recognition helix) by rigid-body fitting, ALC1 residues 267-580 (C-ATPase lobe) by rigid-body fitting, ALC1 residues 28-266 (N-ATPase lobe) by rigid-body fitting. Overlapping segments were de-duplicated by removing one of the segments, to ensure all fragments followed each other in sequence, with no duplicated residue numbers or gaps (other than the N- and C-termini that were trimmed because they were not supported by density). All fragments therefore had termini contiguous in sequence, and rigid-body fitting brought them in close spatial proximity, giving us confidence in the rigid-body fitting solutions. The ALC1 fragments were finally assigned the same chain ID, and all fragments were combined into a single atomic model of the ALC1-nucleosome complex.

Flexible fitting was performed by interactive molecular dynamics in ISOLDE version 1.10.1 (Croll, 2018), with secondary structure restraints, base-pair restraints and reference-model restraints for torsion angles (with the initial model as reference). Attempting to release reference-model restraints led to a substantial degradation of the model geometry, so all restraints were maintained throughout the flexible fitting.

Final refinement was performed with *phenix.real_space_refine* from the Phenix suite version 1.21.2-5419 (Afonine *et al*., 2018). The refinement strategy was as follows: reference model restraints set to the input model, refine *minimization_global* and *adp*. Validation metrics of the model and model to map fit were calculated with *phenix.validation_cryoem* and *phenix.molprobity*. Q-scores (Pintilie *et al*., 2020) were calculated with the *QScore* extension to ChimeraX. Summary statistics of atomic Q-scores and B-factors were calculated with R version 4.5.0 (R Core Team, 2025).

### 6.3. Conservation analysis

The four sequences of ALC1 orthologues with reviewed status available in Uniprot at the time of this writing (*Homo sapiens*: Q86WJ1, *Mus musculus*: Q9CXF7, *Bos taurus*: Q3B7N1, *Danio rerio*: Q7ZU90) were aligned with MUSCLE version 5.2 (Edgar, 2022). The resulting multiple sequence alignment (MSA) was used to build a profile hidden Markov model (pHMM) using *hmmbuild* from the HMMER suite version 3.4 (Eddy, 2011). This pHMM was used to search for ALC1 orthologues in the reference proteomes of the following model organisms: *Gallus gallus* (UP000000539), *Xenopus laevis* (UP000186698), *Drosophila melanogaster* (UP000000803), *Arabidopsis thaliana* (UP000006548) and *Saccharomyces cerevisiae* (UP000002311). These model organisms were chosen as representatives of vertebrate taxa likely to have an ALC1 orthologue. The proteome of *S. cerevisiae* was used as a negative control in which we do not expect to find an ALC1 orthologue, because this organism doesn’t have any PARP enzyme. The proteome of *A. thaliana* was included as a non-animal but multi-cellular out-group. The search was performed using *hmmsearch* with a filter to report matches with an E-value < 10^-100^. Structure predictions of the top hits from each reference proteome were obtained from AlphaFold-DB when available (Varadi *et al*., 2022), or computed with DeepMind’s AlphaFold3 server otherwise (Abramson *et al*., 2024), and inspected individually. From these structure predictions, plausible ALC1 orthologues were identified based on the following criteria: a general architecture with two ATPase lobes, a linker and a macro domain (in this order from N- to C-ter); the presence of Arg residues in the non-helical regions of the linker, as putative Arg-anchor residues defining the RLS; the presence of an alpha helix about 40-residue long just upstream of the macro domain, as a putative SGRH. These criteria retained sequences from *Gallus gallus* and *Arabidopsis thaliana*, and excluded the sequence from *Xenopus laevis* that did not exhibit a recognizable putative SGRH. Despite having a PARP enzyme, *Drosophila melanogaster* does not seem to have an ALC1 orthologue, as the top hits from the pHMM search against its proteome are the remodelers ISWI and CHD1. The sequences of these two additional plausible ALC1 orthologues were aligned to the initial four sequences to produce a new MSA, using MUSCLE. This final MSA was used to color the SGRH in our structure by sequence conservation, using ChimeraX. The figure of the final MSA was produced with Aliview (Larsson, 2014). The final MSA is provided as supporting data.

## Appendix A. ChimeraX command file to set up a session with the same color code and style as in the figures.

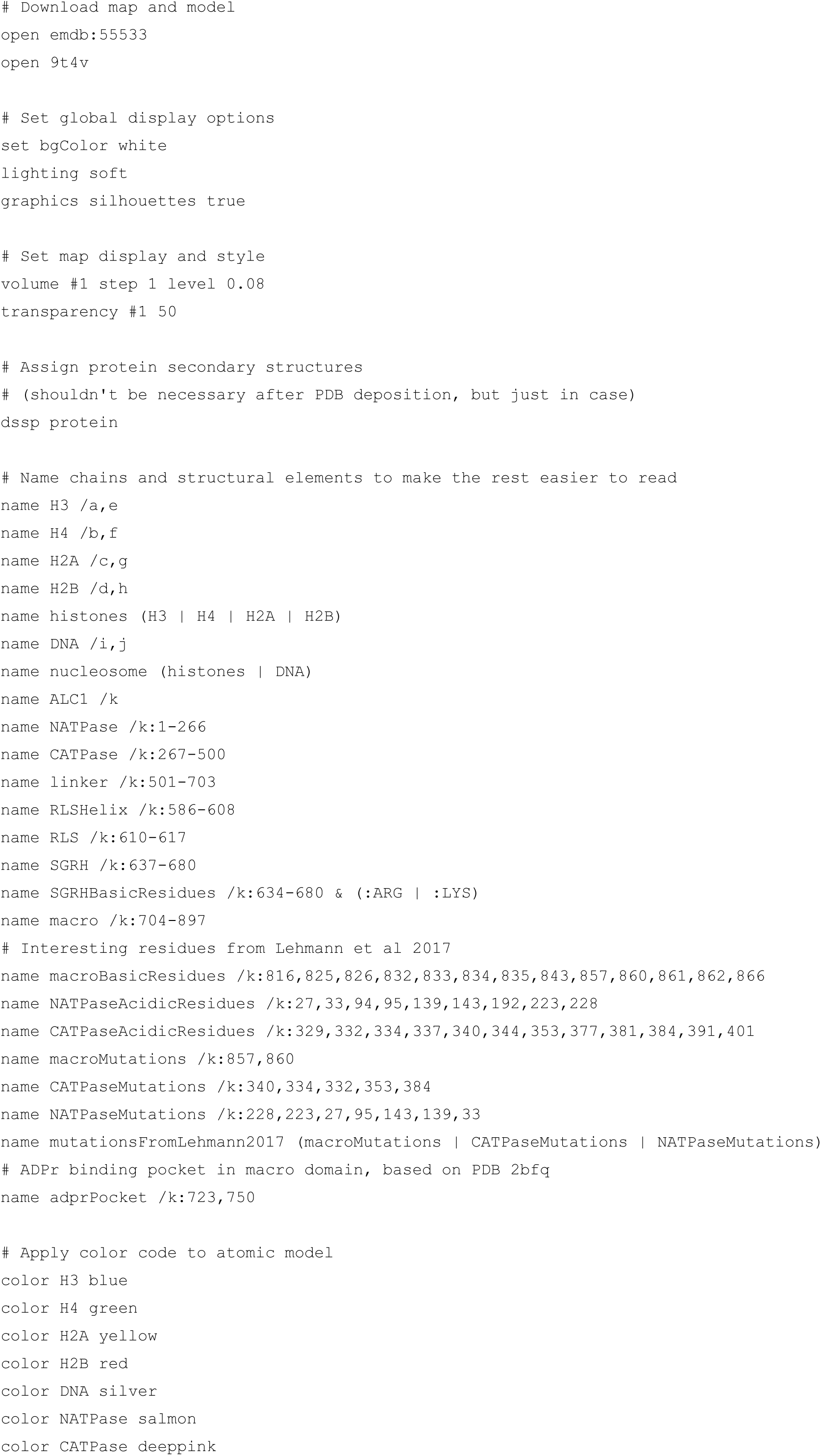
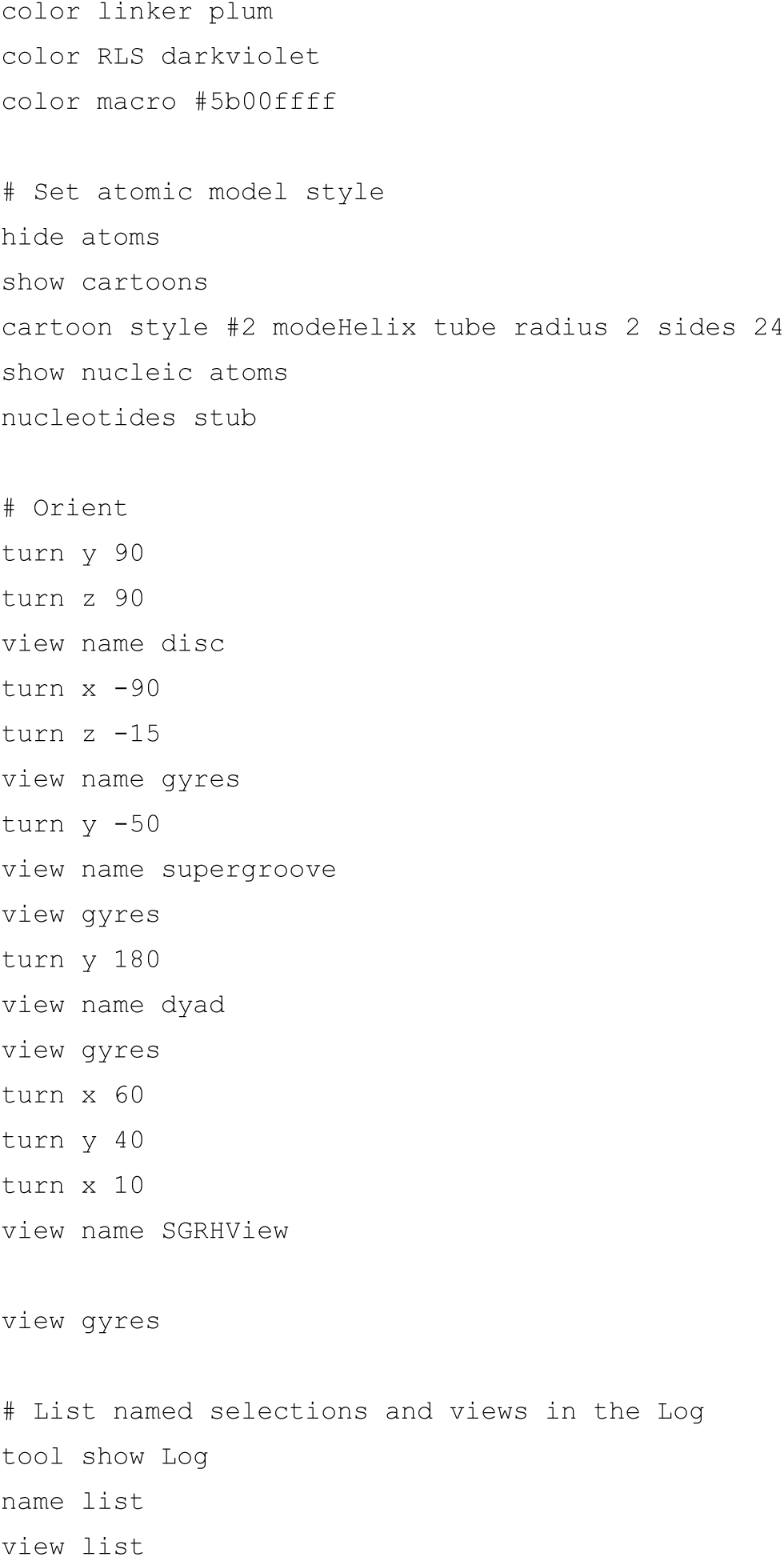

## Supporting information

**Figure S1.**
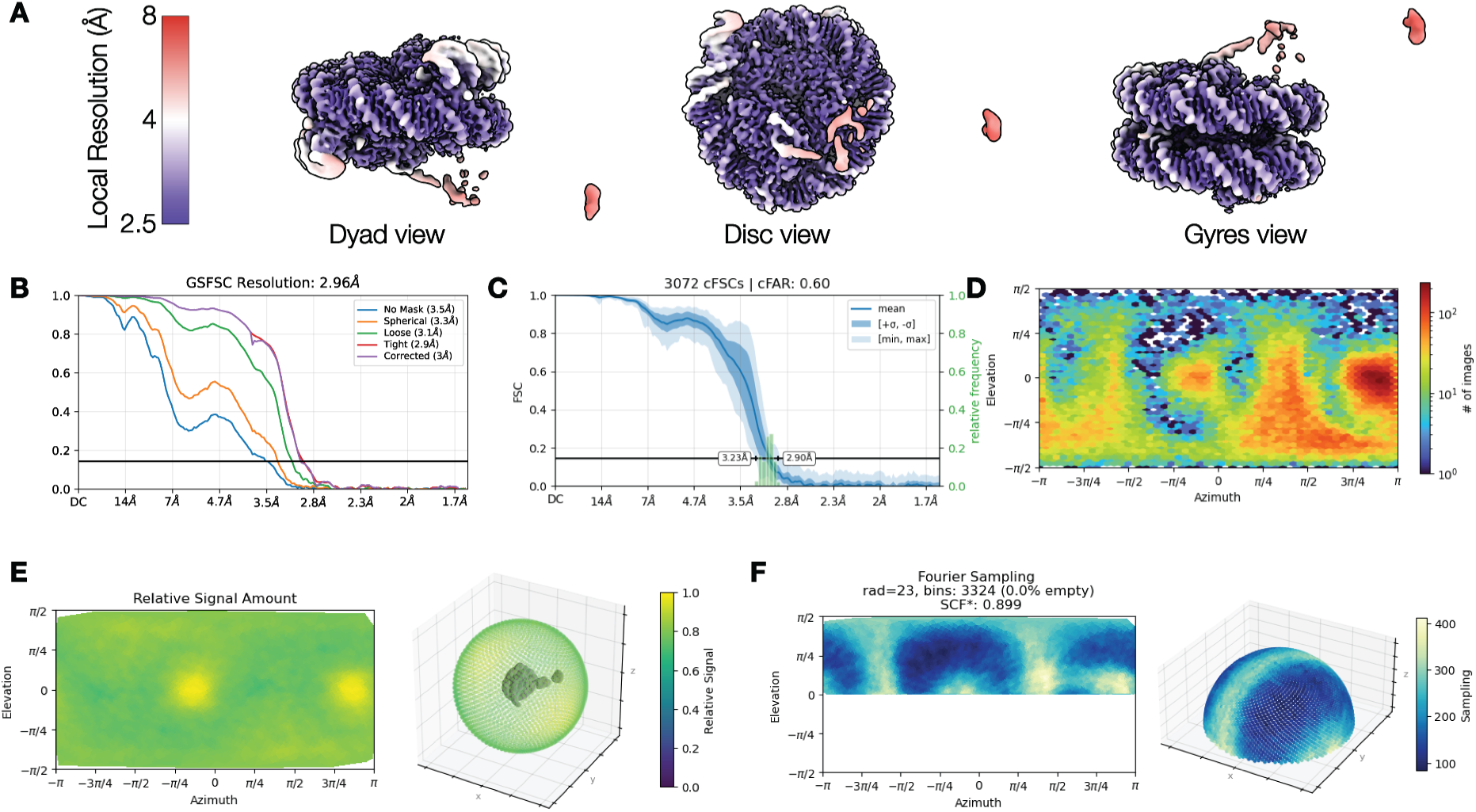
Validation of the cryo-EM map used to build the RLS with ModelAngelo. (A) Map colored by local resolution. (B) Gold-standard FSC curve. (C) Measure of resolution anisotropy by conical FSC area ratio (cFAR). (D) Euler angles distribution of the set of particles contributing to this reconstruction. (E) Relative signal versus viewing direction. (F) Distribution of Fourier sampling.

**Figure S2.**
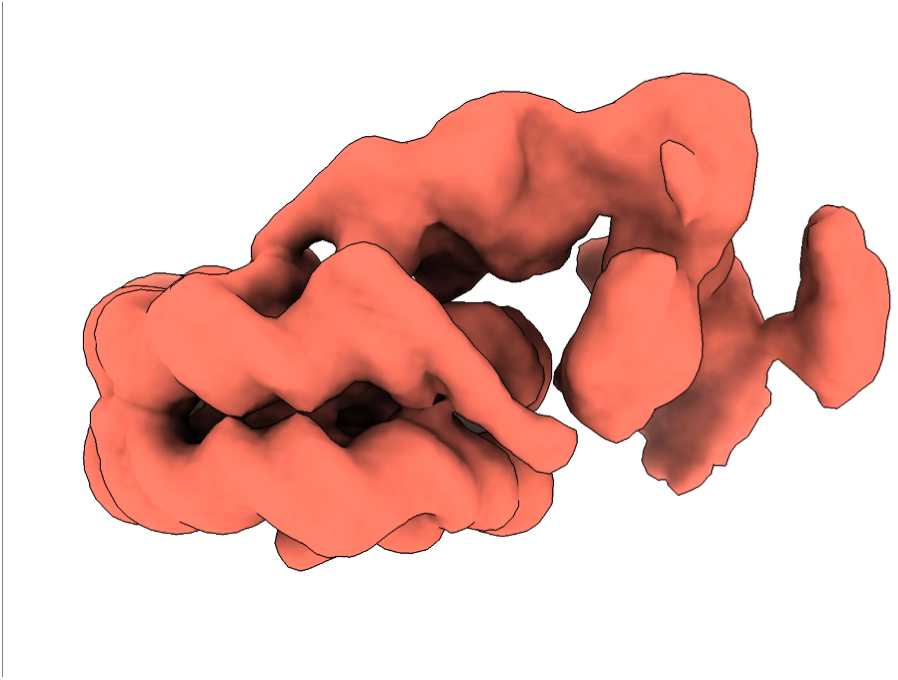
Movie of 3DVA (2).

**Figure S3.**
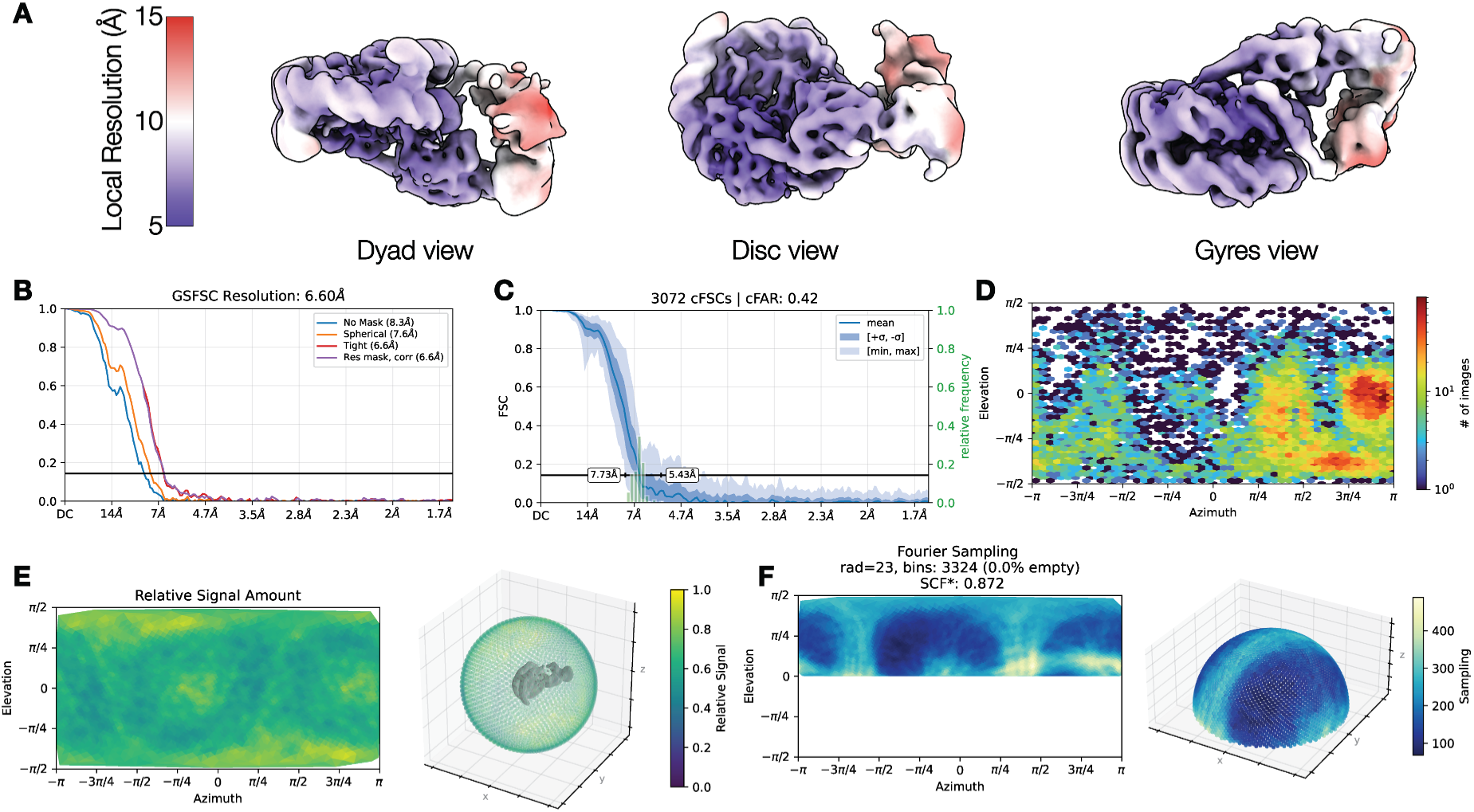
Validation of the main cryo-EM map. (A) Map colored by local resolution. (B) Gold-standard FSC curve. (C) Measure of resolution anisotropy by conical FSC area ratio (cFAR). (D) Euler angles distribution of the set of particles contributing to this reconstruction. (E) Relative signal versus viewing direction. (F) Distribution of Fourier sampling.

**Table S1.** Statistics from data collection, image processing, model building and validation. ND: not determined.

|  |  |  |
| --- | --- | --- |
| **Data collection** | See Table 1 in (Bacic *et al.*, 2021) | See Table 1 in (Bacic *et al.*, 2021) |
| Number of micrographs collected | 33 998 | 33 998 |
| **Image processing** | **RLS resolved (NU Refinement 3)** | **ALC1 activation intermediate (Local Refinement 2)** |
| Number of micrographs used for picking | 28 448 | 28 448 |
| Number of picked particles | 2 001 174 | 2 001 174 |
| Number of particles used for reconstruction | 57 036 | 15 740 |
| Point group symmetry imposed | C1 | C1 |
| Conical FSC area ratio (cFAR) | 0.60 | 0.42 |
| Fourier sampling compensation factor (SCF*) | 0.899 | 0.872 |
| Map sharpening B-factor ($\text{\AA}^2$) | -24.3 | -300 |
| Global resolution at FSC = 0.143 ($\text{\AA}$) | 2.96 | 6.60 |
| **EMDB accession code** | EMD-55534 | EMD-55533 |
| **Model building and refinement** | **RLS resolved (NU Refinement 3)** | **ALC1 activation intermediate (Local Refinement 2)** |
| Initial models | Output of ModelAngelo run with sequence information. | Nucleosome from PDB 8B0A.<br>ALC1 residues 599-626 from the output of ModelAngelo run on the map from NU Refinement 3 with sequence information.<br>Other ALC1 domains from AlphaFold-DB AF-Q86WJ1-F1-v4. |
| Number of protein residues | 796 | 1610 |
| Number of protein atoms (including hydrogen atoms) | 6 320 (ND) | 12 805 (25 971) |
| Number of DNA residues | 312 | 298 |
| Number of DNA atoms (including hydrogen atoms) | 6 377 (ND) | 6 109 (9 454) |
| Number of ligands | 0 | 0 |
| Number of ions | 0 | 0 |
| Number of water molecules | 0 | 0 |
| **Model validation** | **RLS resolved (NU Refinement 3)** | **ALC1 activation intermediate (Local Refinement 2)** |
| Protein B-factors (min/median/max, $\text{\AA}^2$) | ND | 14 / 306 / 903 |
| DNA B-factors (min/median/max, $\text{\AA}^2$) | ND | 67.3 / 319 / 1030 |
| Bond lengths RMSD (Å) | ND | 0.008 |
| Bond angles RMSD (°) | ND | 1.292 |
| MolProbity score | ND | 1.75 |
| Clashscore | ND | 7.48 |
| Rotamer outliers (%) | ND | 0.73 |
| C-beta deviations | ND | 0 |
| Ramachandran favored (%) | ND | 95.16 |
| Ramachandran allowed (%) | ND | 3.52 |
| Ramachandran outliers (%) | ND | 1.32 |
| **Model to map fit** | **RLS resolved (NU Refinement 3)** | **ALC1 activation intermediate (Local Refinement 2)** |
| CC volume | ND | 0.7191 |
| CC mask | ND | 0.7439 |
| CC peaks | ND | 0.5931 |
| Protein Q-scores (min/median/max) | ND | -0.845 / 0.282 / 0.847 |
| DNA Q-scores (min/median/max) | ND | -0.610 / 0.398 / 0.805 |
| **PDB accession code** | Not deposited | pdb_00009T4V |

## Notes

https://www.ebi.ac.uk/empiar/EMPIAR-10739

https://guide.cryosparc.com/processing-data/tutorials-and-case-studies/case-study-end-to-end-and-exploratory-processing-of-a-motor-bound-nucleosome-empiar-10739

